# BECLIN1 is essential for intestinal homeostasis

**DOI:** 10.1101/2023.06.14.545007

**Authors:** Sharon Tran, Juliani Juliani, Tiffany J. Harris, Marco Evangelista, Julian Ratcliffe, Sarah L. Ellis, David Baloyan, Camilla M. Reehorst, Rebecca Nightingale, Ian Y. Luk, Laura J. Jenkins, Sonia Ghilas, Marina H. Yakou, Chantelle Inguanti, Chad Johnson, Michael Buchert, James C. Lee, Peter De Cruz, Kinga Duszyc, Paul A. Gleeson, Benjamin T. Kile, Lisa A. Mielke, Alpha S. Yap, John M. Mariadason, W. Douglas Fairlie, Erinna F. Lee

## Abstract

BECLIN1 is a component of Class III phosphatidylinositol 3-kinase complexes that orchestrates autophagy initiation and endocytic trafficking. Here we show intestinal epithelium-specific BECLIN1 deletion in adult mice led to rapid fatal enteritis with compromised gut barrier integrity, highlighting its intrinsic critical role in gut maintenance. BECLIN1-deficient intestinal epithelial cells exhibited extensive apoptosis, impaired autophagy, and stressed endoplasmic reticulum and mitochondria. Remaining absorptive enterocytes and secretory cells displayed morphological abnormalities. Deletion of the autophagy regulator, ATG7, failed to elicit similar effects, suggesting novel autophagy-independent functions of BECLIN1 distinct from ATG7. Indeed, organoids derived from BECLIN1 KO mice showed E-cadherin mislocalisation providing a mechanism linking endocytic trafficking mediated by Beclin1 and loss of intestinal barrier integrity. Our findings establish an indispensable role of BECLIN1 in maintaining mammalian intestinal homeostasis and uncover its involvement in endocytic trafficking in this process. Hence, this study has significant implications for our understanding of intestinal pathophysiology.

## Introduction

Autophagy is an evolutionarily conserved process critical for the maintenance of eukaryotic cellular homeostasis. It encapsulates cytoplasmic material within autophagosomes and mediates their delivery to the lysosome for degradation. Critically, genome-wide association studies (GWAS) have identified that polymorphisms in genes that regulate autophagy correlate with a susceptibility to inflammatory bowel disease (IBD), which includes Crohn’s disease and ulcerative colitis ^1^. Subsequent studies aiming to understand this connection between autophagy and intestinal health led to the generation of mice deficient for, or bearing disease variants in, autophagy regulators. These studies revealed autophagy-related roles intrinsic to intestinal epithelial cells (IECs), including the protection against enteric pathogen infections ^2,3^, the homeostatic and secretory capacity of specialised IECs, namely Paneth and goblet cells ^4-10^, and the survival of intestinal stem cells (ISCs) ^11^. Beyond IECs, autophagy also contributes to the survival and function of intestinal stromal cells, including immune cells, where disruptions to these autophagic functions have adverse impacts on the intestinal cytokine and growth factor milieu ^12,13^. Whilst these are important roles for autophagy regulators in the maintenance of intestinal homeostasis, autophagy-deficient intestines generally function normally, and autophagy deficiency alone is insufficient for driving spontaneous intestinal pathologies, consistent with the paradigm that IBD pathogenesis is multifactorial.

BECLIN1 is a regulator of autophagy that exerts its cellular functions as a scaffolding subunit of two mutually exclusive Class III phosphatidylinositol 3-kinase (PI3KC3) complexes ^14^. These complexes share two common subunits in addition to BECLIN1: the lipid kinase, Class III phosphoinositide 3-kinase (PI3KC3), and the regulatory subunit, VPS15. Distinguishing the two complexes are autophagy-related protein 14 (ATG14) in Complex 1 (PI3KC3-C1) and UV radiation resistance gene product (UVRAG) in Complex 2 (PI3KC3-C2) ^15-18^. Both complexes promote autophagosome formation and maturation, although notably, PI3KC3-C2 is additionally localised to endosomes and, therefore, can also regulate endocytic trafficking ^16,17^. There is some indication that BECLIN1 may be relevant to intestinal health. For example, using a cell culture model, BECLIN1 was shown to play a constitutive autophagy-independent role in intestinal tight junction barrier function through the endosomal degradation of occludin ^19^. In *Drosophila* ISCs, the orthologue of BECLIN1, ATG6, is required for maintaining the integrity of this cellular compartment in response to aging ^20^. Recently, it was found that mice expressing constitutively active mutant BECLIN1 have a thicker colonic mucosal layer due to autophagy-mediated alleviation of endoplasmic reticulum (ER) stress, demonstrating a potential role for BECLIN1 in the maintenance of intestinal homeostasis ^21^. However, despite these indicators, and the plethora of genetic models in which other key autophagy regulators have been deleted in the intestinal epithelium ^22^, to our knowledge, there is no published murine model in which BECLIN1 has been genetically deleted in this compartment.

In this study, we report for the first time that severe fatal intestinal disruption ensues following intestinal epithelial-specific deletion of BECLIN1 in adult mice. This phenotype is not seen when the autophagy regulator ATG7 is deleted. We postulate that the intestinal architectural disruption, extensive IEC death, and functional abnormalities, leading to the fatal crippling of intestinal homeostasis following BECLIN1 loss also involves autophagy-independent mechanisms, likely caused by defective endocytic trafficking.

## Results

### Generation of conditional *Becn1* and *Atg7* intestinal epithelium-specific inducible knock-out mice

To induce intestinal epithelium-specific deletion of *Becn1*, *Becn1^fl/fl^* mice were bred to mice expressing tamoxifen-inducible Cre recombinase from the villin 1 promoter (*Vil1-CreERT2,* Fig. 1a). For comparison, we also generated mice in which the pro-autophagic E1-like ligase ATG7, which unlike Beclin1 has no reported roles in endocytic trafficking ^23^, could be deleted in a similar manner (Fig. 1a). The resulting tamoxifen-treated *Becn1^fl/fl^*;*Vil1-CreERT2^Cre/+^* and *Atg7^fl/fl^*;*Vil1-CreERT2^Cre/+^*mice will hereafter be referred to as *Becn1^ΔIEC^* and *Atg7^11IEC^* mice respectively, whilst their littermate tamoxifen-treated *Becn1^+/+^*;*Vil1-CreERT2^Cre/+^*and *Atg7^+/+^*;*Vil1-CreERT2^Cre/+^* mice are referred to as *Becn1^wtIEC^* and *Atg7^wtIEC^* mice respectively (Fig. 1a). All time points, unless otherwise specified, indicate time after receipt of the first tamoxifen dose. Inducible conditional deletion of either *Becn1* or *Atg7* in the intestinal epithelium was confirmed by genotyping (Supp. Fig. 1) and Western blotting (Fig. 1b) of IECs isolated from *Becn1^ΔIEC^* and *Atg7^11IEC^* mice. Both showed significantly reduced BECLIN1 and ATG7 expression, respectively, compared to control wild-type mice after seven days. As expected, loss of either BECLIN1 or ATG7 resulted in disrupted basal autophagic flux as indicated by an accumulation of p62 and LC3 protein levels, or an increased LC3-I:LC3-II ratio, in *Becn1^ΔIEC^*- and *Atg7^11IEC^*-derived IECs (Fig. 1b). Our successful generation of mice in which either BECLIN1 or ATG7 can be specifically deleted in the intestinal epithelium in an inducible manner, enabled us to investigate the role of BECLIN1 in this tissue. In addition, the comparison with ATG7 deleted mice enabled delineation of the contributions of autophagy *versus* endocytic trafficking mechanisms to this process.

**Fig. 1.**
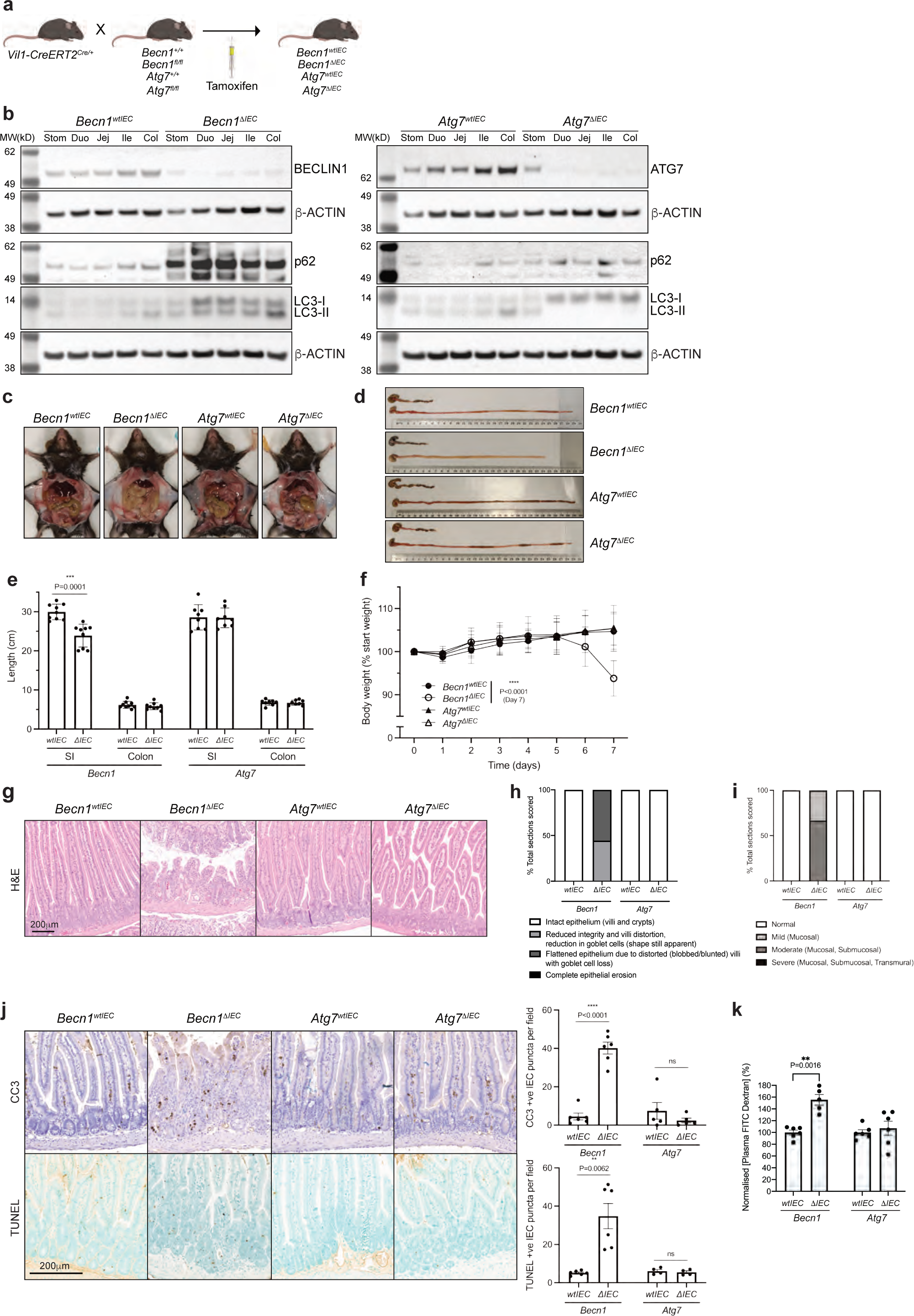
The absence of BECLIN1, but not ATG7, leads to fatal loss of intestinal homeostasis. **a** Intestinal-epithelium specific deletion of loxP-flanked *Becn1* or *Atg7* in adult mice is mediated by tamoxifen-inducible Cre recombinase from the villin 1 promoter. **b** Deletion of either BECLIN1 or ATG7 leads to defective autophagy as indicated by increased levels of total p62 and LC3, or increased LC3-I relative to LC3-II. Representative images of **c** abdominal necropsy and **d** intestinal tracts demonstrating the severe intestinal disruption seen only in the absence of BECLIN1 (upper: colon; lower: small intestine). **e** Intestinal lengths of mice at endpoint. **f** Body weight of mice over time normalised to day 0. **g** FFPE sections of mouse intestines, stained with H&E, reveal the extensive loss of the characteristic villus-crypt structures of the small intestines from *Becn1^11IEC^*mice. **h** Scoring of intestinal morphological integrity**. i** Loss of intestinal homeostasis in the absence of BECLIN but not ATG7 leads to increased mucosal and/or submucosal immune cell infiltration. **j** There was evidence of increased apoptosis, as determined by cleaved caspase-3 (CC3) and terminal doxynucleotidyl transferase dUTP nick-end labelling (TUNEL) staining, along the crypt-villus axis in *Becn1^11IEC^* mice only. **k** Loss of BECLIN1 results in loss of intestinal barrier integrity leading to increased permeability to FITC-Dextran. All data represent eight mice of each genotype, except for the histology images which represent a minimum of four mice per genotype. Data represent 3 independent experiments unless otherwise indicated. Graphs show the mean ± S.E.M, except for the small intestinal lengths and body weight graphs which show the mean ± S.D. Significance was determined by Welch’s unpaired test. ns = not significant (P > 0.05). Stom: stomach. Duo: duodenum. Jej: jejunum. Ile: ileum. Col: colon. SI: small intestine.

### Deletion of BECLIN1, but not ATG7, in the intestinal epithelium, leads to severe fatal loss of intestinal homeostasis

Strikingly, within seven days post-induction, *Becn1^11IEC^* mice developed severe and fatal enteritis with 100% penetrance with the small intestine appearing lytic, necrotic and grossly swollen (Fig. 1c, d). These mice presented with shortened small intestinal, but not colon, lengths (Fig. 1d, e) and significant body weight loss (>10%, Fig. 1f), with humane euthanasia being required. Inspection of all other organs, including the colon, at this time point, revealed they were grossly anatomically normal and similar to their wild-type littermates. Notably, this phenotype contrasted with *Atg7^11IEC^* mice over the same time course (Fig. 1c-f); these mice did not develop any fatal or macroscopic intestinal pathologies when the same parameters were assessed, even when aged up to one month (Supp. Fig. 2). However, interestingly, at this time point, they displayed a small but significant decreased weight gain and increased intestinal length compared to wild-type controls (Supp. Fig. 2).

**Fig. 2.**
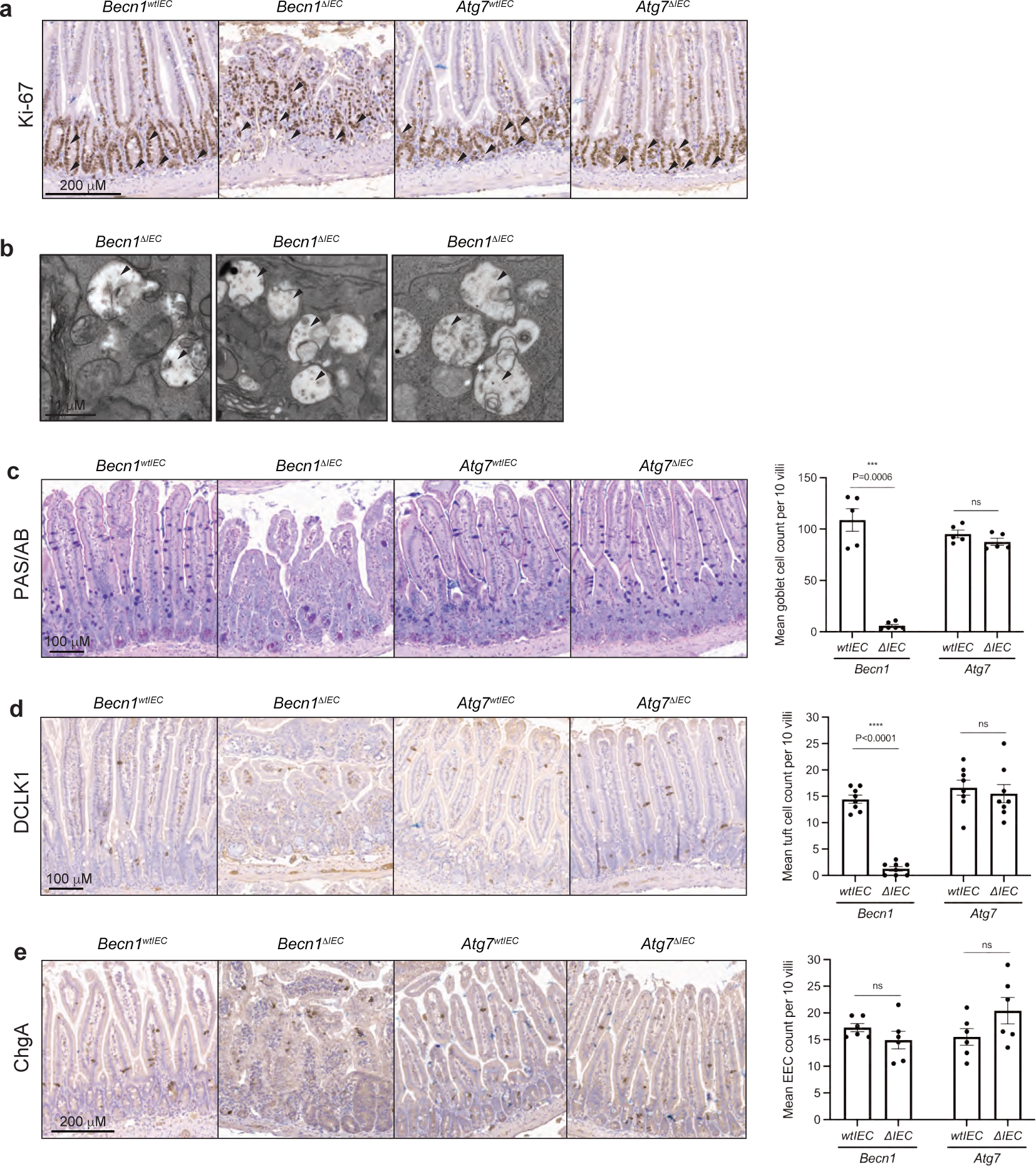
BECLIN1 deletion leads to loss of differentiated intestinal epithelial cell sub-populations. **a** The proliferative capacity of cells in the intestinal stem cell compartment and transit amplifying regions (black arrows), remains intact when BECLIN1 or ATG7 is absent. **b** However, there is evidence of abnormal multivesicular body (black arrows) accumulation, suggesting compromised endocytic trafficking, in BECLIN1-deficient transit-amplifying cells. FFPE sections of small intestines were stained with specific markers for the various IEC sub-populations. BECLIN1 is critical for the maintenance of specific IEC sub-populations such as **c** goblet (Periodic acid-Schiff-Alcian Blue, PAS/AB staining) and **d** tuft (doublecortin like kinase 1, DCLK1, staining) cells, but not others such as **e** enteroendocrine (chromogranin A, ChgA staining) cells. Quantitation of these specific IECs is represented in graphs on the right. All data represent a minimum of at least four mice from three independent experiments. Graph shows the mean ± S.E.M. Significance determined by Welch’s unpaired t-test. ns = not significant (P > 0.05). EEC: enteroendocrine cells.

Characterisation of the intestinal pathology of *Becn1^11IEC^* mice by haematoxylin and eosin (H&E) staining revealed dramatic architectural changes, which included the loss of intestinal morphological integrity with villi blunting (scored using Erben *et al.* classifications ^24^ (Fig. 1g, h), and evidence of mucosal and submucosal, though no transmural, immune infiltration (Fig. 1g, i). Staining of intestinal sections with cleaved caspase-3 (CC3) and terminal deoxynucleotidyl transferase dUTP nick-end labelling (TUNEL) showed widespread apoptosis of epithelial cells both in the crypt and along the villi length (Fig. 1j). Flow cytometry-based immunophenotyping of intraepithelial lymphocytes (IELs) isolated from these mice showed an increased proportion of total lymphocytes, specifically T cells (Supp Fig. 3). In particular, there was a statistically significant increase in the numbers of cytotoxic CD8+ T cells. There was also a trend towards increased gamma-delta T cells important for immune surveillance at epithelial and mucosal surfaces, though this was not statistically significant. Consistent with the dramatic morphological disruption and loss of IEC survival in *Becn1^11IEC^* mice, we observed a severely compromised total intestinal barrier as determined by the increased release of fluorescein isothiocyanate-dextran (FITC-dextran) from the intestinal lumen into the circulation of these mice (Fig. 1k). Importantly, these dramatic intestinal disruptions were not observed in *Atg7^11IEC^* mice over the same time frame (Fig. 1g-k, Supp. Fig. 3).

**Fig. 3.**
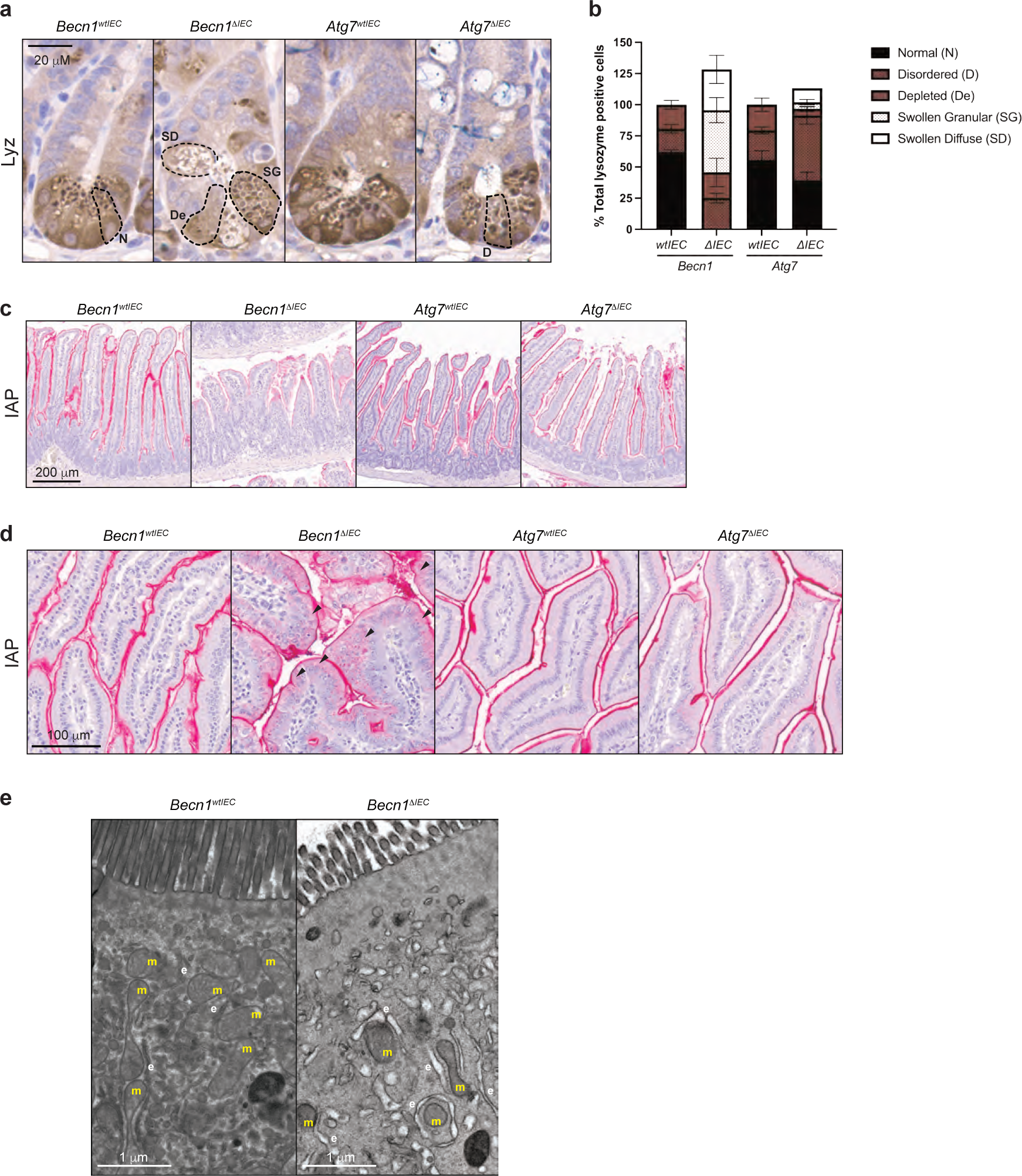
BECLIN1 is critical for maintaining the specialised functions of secretory intestinal epithelial cells. FFPE sections of small intestines were stained for the anti-microbial proteins lysozyme (Lyz) and intestinal alkaline phosphatase (IAP) produced by both Paneth and enterocytes, respectively. **a** Dashed lines delineating the borders of Paneth cells demonstrate the significant morphological abnormalities of lysozyme-containing secretory granules from both *Becn1^11IEC^*and *Atg7^11IEC^* animals, though the defects in the absence of BECLIN1 are far more pronounced. **b** Graphical representation of the morphological classifications of abnormal secretory granules in Paneth cells. Villus-associated enterocytes deficient for BECLIN1 demonstrate **c** almost absent IAP expression or where IAP is present, **d** diffuse cytoplasmic staining (black arrows) as opposed to well-defined membrane localisation. **e** Transmission electron microscopy (TEM) images of *Becn1^11IEC^* and *Becn1^wtIEC^* enterocytes reveal distorted mitochondria (m) and electron-lucent tubulated structures characteristic of swollen ER (e) and ER stress in the former. All data represent a minimum of at least four mice from three independent experiments. In the case of the TEM analysis, data shown are representative of two independent experiments. Graph shows the mean ± S.D.

Taken together, our results demonstrate an essential role for BECLIN1 in the maintenance of intestinal homeostasis. We postulate that its autophagy-independent function in endocytic trafficking likely contributes significantly given the striking intestinal phenotypic differences between *Becn1^ΔIEC^* and *Atg7^11IEC^* mice.

### BECLIN1 is essential to the survival and function of specialised intestinal epithelial cells

The intestinal epithelium is made up of a single layer of cells comprising differentiated IEC subpopulations, all of which arise from ISC progenitors in the crypt base. Each IEC subpopulation confers their own distinct specialised function, which together regulate intestinal homeostasis ^25^. Staining for the proliferation marker Ki-67 in *Becn1^11IEC^* crypts and transit-amplifying regions was retained on day 7, as in *Atg7^11IEC^* and control mice (Fig. 2a), indicating that the proliferative capacities of ISCs and transit-amplifying progenitor cells (TACs) are not adversely impacted by BECLIN or ATG7 loss at this time point. Interestingly, ultrastructural analysis by transmission electron microscopy (TEM) of the intestinal crypts of *Becn1^11IEC^* mice revealed enlarged single-membrane electron lucent structures containing heterogenous contents, in the transit-amplifying region (Fig. 2b). This is suggestive of abnormal multivesicular body accumulation, implicating defective endocytic trafficking within BECLIN1-deficient transit-amplifying cells.

To interrogate any IEC-specific changes within the intestinal epithelium following BECLIN1 or ATG7 deletion on day 7, we assessed each of the major IEC subtypes in our knock-out mice using immunohistological techniques. We observed a significant loss of specific IEC subpopulations, in *Becn1^ΔIEC^* but not *Atg7^11IEC^* or wild-type littermates. Specifically, there were significant reductions in tuft (DCLK1^+ve^) and goblet cell (PAS/AB^+ve^) numbers (Fig. 2c, d). In contrast, there were no significant changes to the numbers of enteroendocrine (ChgA^+ve^, Fig. 2e) or Paneth subpopulations.

Of the IEC subpopulations that remained significantly unchanged in numbers, there were instead observable abnormalities in their morphologies suggestive of functional disruptions pertaining to their anti-microbial capacities. Specifically, inspection of *Becn1^11IEC^* Paneth cells revealed drastic morphological changes, with an increased proportion of cells being “depleted” (using the lysozyme staining pattern allocations published by Cadwell *et al*. ^6^), and containing fewer granules and reduced lysozyme levels compared to wild-type mice (Fig. 3a, b). Other Paneth cells appeared swollen with aberrantly accumulated lysozyme-positive granules, which we have newly assigned as “swollen granular” (Fig. 3a, b). Some swollen cells also displayed a “ruptured” appearance, which we have further categorised as “swollen diffuse” (Fig. 3a, b). Whilst Paneth cells from *Atg7^11IEC^* mice also demonstrated altered granule integrity with a shift to a more disordered patterning which is consistent with published reports (Fig. 3a, b) ^8,26^, these defects were not as extensive as those observed in *Becn1^11IEC^* mice.

In addition to Paneth cells, we also observed defects in staining for intestinal alkaline phosphatase (IAP), a component of the gut mucosal defense system normally expressed at the brush border of villus-associated enterocytes. Levels of IAP were visibly reduced, particularly towards the distal end of the small intestine, in *Becn1^ΔIEC^* but not *Atg7^11IEC^* or wild-type mice (Fig. 3c). Furthermore, even though staining in the duodenum at the proximal end of the small intestine was still evident, there was noticeable mislocalisation of IAP, shifting from a well-defined border staining to a more diffuse cytoplasmic distribution (Fig. 3d). Ultrastructural examination of *Becn1^11IEC^* enterocytes by TEM found electron lucent tubulated single membrane structures resembling a swollen endoplasmic reticulum (ER) compartment, suggestive of ER stress (Fig. 3e). There was also evidence of degenerating mitochondria (Fig. 3e) and complete disappearance of autophagosomes as expected.

Taken together, these results indicate, for the first time, significant roles for BECLIN1 in maintaining the survival and anti-microbial functions of the different specialised epithelial cells that constitute the barrier function of the intestinal epithelium. Importantly, its coordinated regulation of intestinal homeostasis goes far beyond that implicated for the autophagy regulator ATG7 ^23^.

### BECLIN1 deletion in small intestinal organoids leads to increased apoptosis and morphological defects

Our results indicate essential IEC-intrinsic contributions made by BECLIN1 to the heterogenous, differentiated cell-types of the small intestine and for the maintenance of intestinal integrity and homeostasis. In order to further probe the mechanisms by which BECLIN1 contributes to these, and given the IEC-intrinsic nature of these functions, we isolated and expanded intestinal crypts from *Becn1^fl/fl^*; *Becn1^+/+^*; *Atg7^fl/fl^*; and *Atg7^+/+^*;*Vil1-CreERT2^Cre/+^*mice to generate intestinal epithelial organoid cultures. Treatment with the tamoxifen metabolite, 4-hydroxytamoxifen (4-HT) induced deletion of BECLIN1 or ATG7 in *Becn1^fl/fl^*;*Vil1-CreERT2^Cre/+^* (*Becn1^ΔIEC^*) and *Atg7^fl/fl^*;*Vil1-CreERT2^Cre/+^*(*Atg7^11IEC^*) respectively but not in *Becn1^+/+^*;*Vil1-CreERT2^Cre/+^* (*Becn1^wtIEC^*) or *Atg7^+/+^*;*Vil1-CreERT2^Cre/+^*(*Atg7^wtIEC^*) organoids (Supp. Fig. 4).

**Fig. 4.**
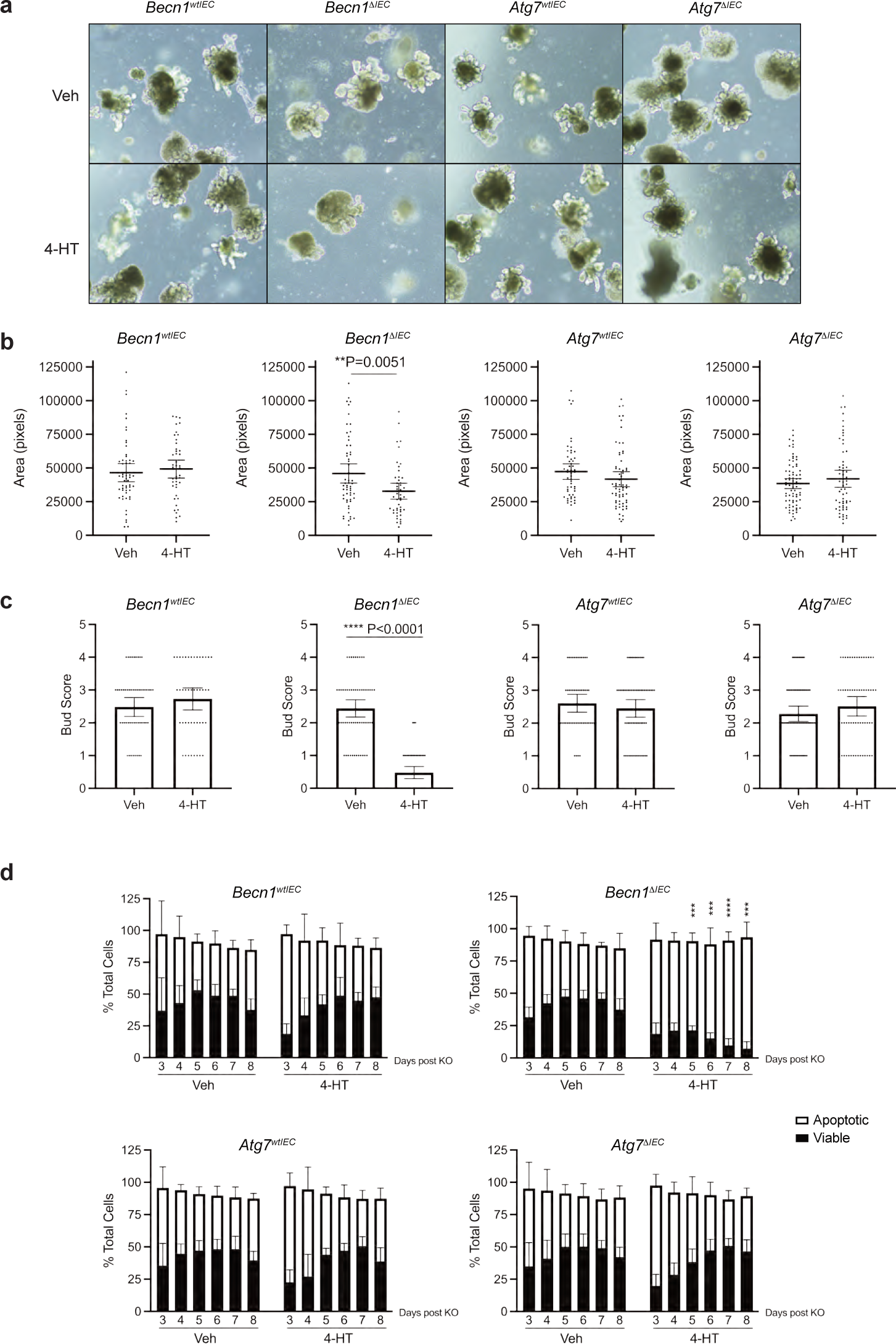
Loss of BECLIN1 in intestinal organoids display increased apoptosis and morphological defects. **a, b** Deletion of BECLIN1, but not ATG7, by addition of 4-hydroxytamoxifen (4-HT) led to significantly smaller intestinal organoids. **a, c** There was also a significant reduction in the number of “buds” per organoid formed, indicative of reduced stem cell-containing crypt formation. **d** BECLIN1- and ATG-deficient organoids were analysed for propidium iodide (PI) and Annexin V staining by flow cytometry, to detect apoptotic cell death, over days 3 to 8 post the first 4-HT dose. Consistent with our *in vivo* observations, we saw enhanced apoptosis in the *Becn1^11IEC^* but not *Atg7^11IEC^* or wild-type organoids. P=0.000573 (day 5), P=0.000277 (day 6), P<0.0001 (day 7), P=0.000754 (day 8). Data are representative of at least three independent experiments. Graphs indicate the mean ± 95 % confidence interval, except for the FACS plot in **d** which indicate mean ± S.D. Significance determined by Welch’s unpaired t-test.

Similar to the effects seen in mouse intestine, eight days following the first dose of 4-HT treatment, *Becn1^11IEC^* organoids were significantly smaller and displayed reduced numbers of budding crypt-like domains per organoid compared to its vehicle-treated wild-type equivalent (Fig. 4a-c). In contrast, *Atg7^11IEC^*organoids did not display the morphological defects seen in the absence of BECLIN1 and were similar in appearance to *Becn1^wtIEC^* and *Atg7^wtIEC^* organoids (Fig. 4a-c). Consistent with the extensive IEC apoptosis, and time to death, seen in the intestinal epithelia of *Becn1^ΔIEC^* mice, we also observed exacerbated apoptosis from day 5 in *Becn1^11IEC^* organoids as determined by Annexin V/Propidium Iodide (PI) staining by flow cytometry analysis (Fig. 4d). In contrast, we did not see the same extent of apoptotic death in *Atg7^11IEC^*, *Becn1^wtIEC^*and *Atg7^wtIEC^* organoids (Fig. 4d). Notably, the increased apoptosis observed over time in intestinal organoids following BECLIN1 deletion is consistent with the timing of IEC apoptotic death and manifestation of the fatal intestinal phenotype of *Becn1^11IEC^* mice.

### BECLIN1 loss leads to mislocalisation of E-cadherin

Thus far, our data show that BECLIN1 loss leads to a compromised intestinal barrier. To provide a functional link between the increased intestinal permeability seen in *Becn1^11IEC^* mice, and the role of BECLIN1 in endocytic trafficking, we focused on E-cadherin, a major constituent of the adherens junction which mediates cell-cell adhesion. Correct cellular localisation of E-cadherin involves several steps of endocytic trafficking, including Rab5^+ve^ early endosomes ^27^ and Rab11^+ve^ recycling endosomes ^28,29^. These trafficking vesicles themselves have been shown to be regulated by BECLIN1 ^30,31^. Stable E-cadherin membrane localisation is critical for epithelial barrier integrity, and its loss leads to fatal intestinal disruption in mice ^32^, strikingly reminiscent of our *Becn1^11IEC^* phenotype. Recently, BECLIN1 was also shown to mediate E-cadherin surface localisation to adherens junctions in breast cancer cells ^33^. Due to these multiple connections, we performed whole-mount immunofluorescent staining of E-cadherin in intestinal organoids to gain insights into how BECLIN1 deletion impacts intestinal homeostasis. Whilst *Becn1^wtIEC^* organoids displayed the expected continuous linear pattern along the lateral cell borders, and predominant localisation at the apical surface of E-cadherin, its localisation in *Becn1^11IEC^* organoids was significantly different (Fig. 5a). Here, the prominent and well-defined linear staining along both the lateral borders and apical surface of the intestinal epithelial cells was absent (Fig. 5a). Instead, punctate cytoplasmic E-cadherin staining at the apical end of these cells was observed (Fig. 5a). Notably, ATG7 deletion did not impact E-cadherin localisation and neither BECLIN1 nor ATG7 loss altered total cellular E-cadherin levels (Fig. 5b). As such, the mislocalisation of E-cadherin, accompanied by compromised intestinal barrier integrity, in the absence of BECLIN1 but not ATG7, highlights the contribution of BECLIN1-mediated endocytic trafficking in maintaining intestinal homeostasis.

**Fig. 5.**
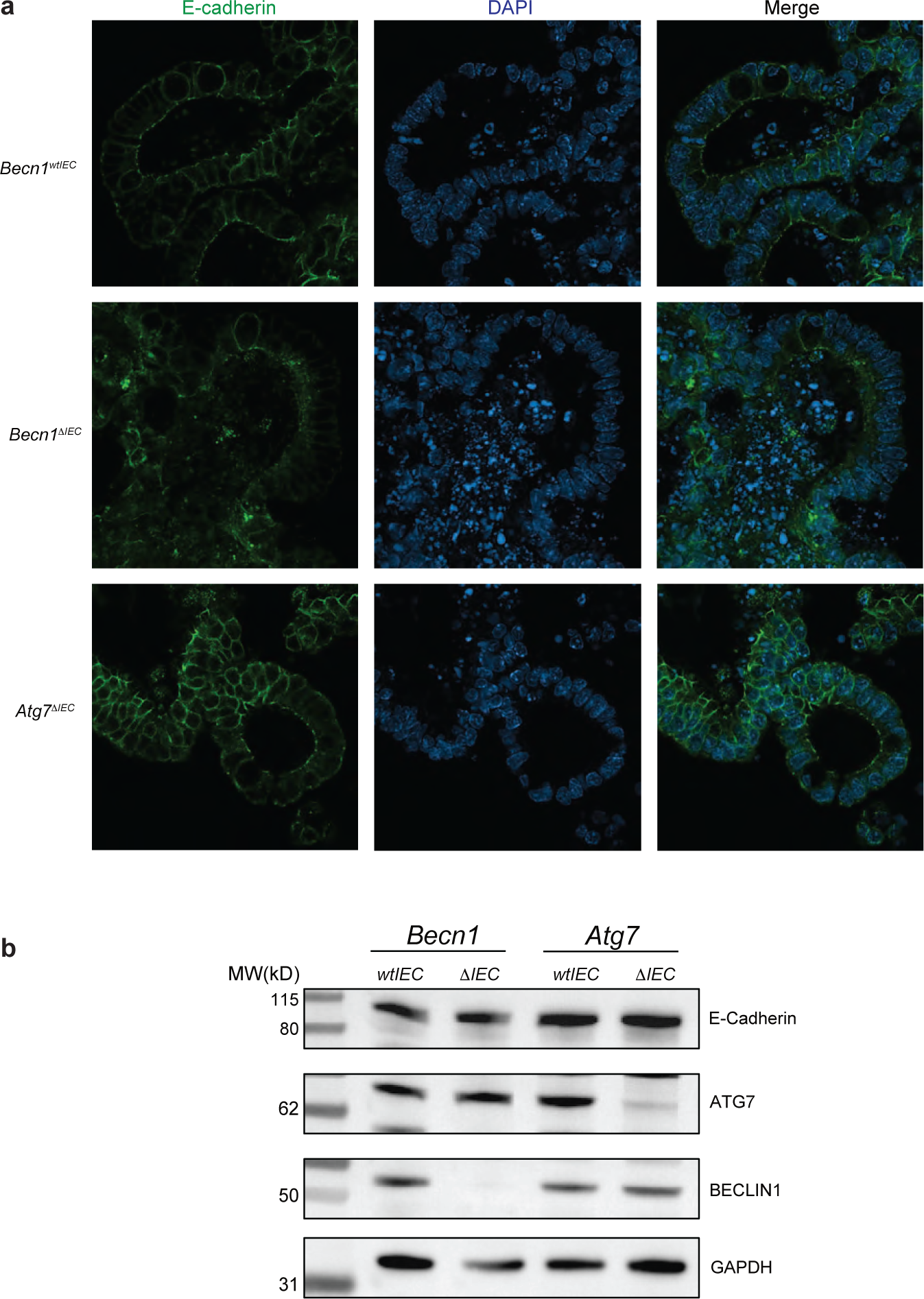
BECLIN1 regulates the correct localisation, but not total cellular levels, of E-cadherin in intestinal epithelial cells. **a** The absence of BECLIN1, but not ATG7, leads to loss of lateral and apical membrane staining of E-cadherin as detected by whole-mount immunofluorescent staining of intestinal organoids. This is consistent with the increased intestinal permeability phenotype observed in *Becn1^11IEC^* mice. **b** Notably, it is the cellular localisation and not the levels of E-cadherin that is altered when BECLIN1 is absent. Data are representative of at least three independent experiments.

## DISCUSSION

We have shown that intestinal epithelial-specific BECLIN1 loss in adult mice results in their rapid and fatal deterioration due to an intestinal pathology marked by intestinal epithelial cell death, inflammation and loss of barrier integrity. For the first time, we demonstrate that BECLIN1 is critically required for the survival and specialised functions of multiple IEC subtypes. These include the maintenance of tuft cells and goblet cells, the regulation of secretory granules in Paneth cells, and the expression and localisation of intestinal alkaline phosphatase in enterocytes. Together, these IEC defects, brought about by the loss of BECLIN1, compromise the intestinal barrier function and overall defence mechanisms of the gut. The significant divergence of phenotypes between *Becn1* loss, and loss of the autophagy regulator *Atg7* in the intestinal epithelium, also led us to postulate that the role of BECLIN1 in endocytic trafficking, which ATG7 is not known to be involved in ^23^, may at least be in part driving these critical functions in the IECs. Notably, we show that in contrast to ATG7, BECLIN1 plays a pivotal role in maintaining appropriate E-cadherin distribution and preserving intestinal permeability. Indeed, loss of cell-cell adhesion *via* the adherens junction has previously been shown to be critical for epithelial barrier integrity in the gut. Taken together, our data now convincingly provide the first known indicators of a physiologically significant role for BECLIN1 in a mammalian intestinal system.

The connection between endocytic trafficking and intestinal physiology has not been widely researched. Our data here characterising *Atg7* intestinal epithelial-specific knock-out mice are consistent with published roles of autophagy in the intestinal epithelium, where a loss of autophagy conjugation proteins including ATG5, ATG16 and ATG7, results in Paneth and goblet cell defects, minimally affects enterocytes, and mice do not develop enteritis unless a chemical or pathogenic agent is administered ^1,22^. Interestingly, we did not observe apoptosis and loss of intestinal stem cells in our ATG7-deficient mice, as reported in Trentesaux et al. ^11^, but this is potentially due to our assessment of mice at a much earlier time point. However, whilst overlapping features are apparent when comparing *Atg7*- and *Becn1*-deficient mice in terms of Paneth and goblet cell changes, *Becn1*-deficient mice have dramatic differences suggestive of non-autophagic roles that are consistent with known roles of other PI3KC3 complex members.

Indeed, there is also some data from non-mammalian organisms that suggest endocytic trafficking could play an important role in the intestine. For example, knock-down of *Uvrag* in *Drosophila* ISCs was associated with impaired differentiation, intestinal dysplasia, and reduced lifespan, attributed to endocytic trafficking defects ^34^. Mislocalised IEC E-cadherin and postnatal lethality with IBD-like features were observed in *Pik3c3*-deficient zebrafish ^35^. In mammalian systems, intestinal epithelial loss of either *Atg14* (or *Fip200*) in mice results in a similar villi atrophy and apoptosis phenotype to *Becn1*-deficient mice, however, this took six weeks to manifest ^36^, probably due to the PI3KC3 C1-specific nature of ATG14 whilst BECLIN1 acts in both PI3KC3 C1 and PI3KC3 C2. Endocytic trafficking also contributes to goblet cell degranulation, requiring endolysosomal convergence with autophagosomes ^5^. In enterocytes, the recycling endosome proteins RAB8 and RAB11 are necessary for the localisation of apical peptidases, transporters and toll-like receptors (TLR), where loss results in intracellular vacuoles, mislocalised proteins, and for *Rab11a* spontaneous enteritis resembling our *Becn1*-deficient mice phenotype ^37,38^. However, differences in phenotype do exist, specifically deficiency in RAB8 or RAB11 is associated with the formation of microvillus inclusion bodies ^37,38^ which is not apparent in *Becn1* deficient mice, suggesting different mechanistic pathways between BECLIN1 and these RAB proteins. However, we did observe evidence of abnormal multivesicular body accumulation, further implicating defective endocytic trafficking in the phenotype of the *Becn1*-deficient mice

The delineation between the autophagy and endocytic trafficking roles of BECLIN1, and their impacts on various IEC subtypes and barrier integrity may yield important knowledge for our understanding of basic intestinal biology and pathophysiology. Indeed, the phenotype of our BECLIN1-deficient mice is strongly reminiscent of the intestinal damage that features in inflammatory bowel disease (IBD). Although BECLIN1, unlike other autophagy regulators, has not previously been linked to IBD, our data reveal that it is a hitherto unknown essential *master regulator* of intestinal homeostasis controlling multiple facets criticial for this process. Hence, further studies examining a potential role for BECLIN1 in IBD are warranted.

## Methods

### Mouse studies

Becn1^tm1b(KOMP)Wtsi^ mice were purchased from the European Conditional Mouse Mutagenesis Program (EUCOMM). *Becn1^fl/fl^* mice were generated by breeding Becn1^tm1b(KOMP)Wtsi^ mice onto CAG-FLPe C57BL/6J mice. These, and the *Atg7^fl/fl^* (Atg7^tm1Tchi^) mice ^39^ were provided by the Kile Laboratory (University of Adelaide, SA, Australia). The *Vil1-CreERT2* mice ^40^ were provided by the Mariadason Laboratory (Olivia Newton-John Cancer Research Institute). Mice were housed at the La Trobe Animal Research and Teaching Facility (LARTF, La Trobe University, VIC, Australia) under Specific Pathogen Free (SPF) conditions. To induce deletion, male and female mice aged six weeks or older were intraperitoneally injected with 4 mg tamoxifen (Sigma-Aldrich, T5648) in sunflower seed oil (Sigma-Aldrich, 25007), delivered as one injection per day of 200 μL of a 10 mg/mL stock, over two consecutive days. Mice were humanely end-pointed by CO_2_ asphyxiation. For PCR genotyping for colony maintenance and KO assessment, DNA extracts were prepared from ear clips or isolated IECs (see IEC isolation) by their overnight incubation at 56°C followed by 1 hour at 85°C in DirectPCR (Tail) Lysis Reagent (Viagen) supplemented with 0.2 % (v/v) Proteinase K (Sigma-Aldrich). PCR reactions were prepared with GoTaq PCR Master Mix (Promega) following the manufacturer’s protocol. Primers for *Becn1* reactions are as follows: “floxed forward” - 5’CTG ATC CTG CAG CTT GCA GAT TAG C3’, “floxed reverse” - 5’CAC CAC TGC CTG GCT AAA CAA GAG C3’, “KO reverse” - 5’CTA TAG AAG AAA GGA CTG TTG TGA AC3’. Primers for *Atg7* reactions are as follows: “floxed forward” - 5’TGG CTG CTA CTT CTG CAA TGA TGT3’, “floxed reverse” -5’CAG GAC AGA GAC CAT CAG CTC CAC3’, “WT forward” – 5’TCT CCC AAG ACA AGA CAG GGT GAA3’, “WT reverse” - 5’AAG CCA AAG GAA ACC AAG GGA GTG3’. Primers for *Vil1-CreERT2* reactions are as follows: forward - 5’CAA GCC TGG CTC GAC GGC C3’, reverse - 5’CGC GAA CAT CTT CAG GTT CT3’. For intestinal barrier permeability experiments, mice were fasted overnight and then orally gavaged with 150 μL of 10 mg/mL FITC-dextran (Sigma-Aldrich, FD4) in DPBS (Gibco). Blood was collected into Microvette® EDTA K tubes (Sarsdtedt) both prior to gavage *via* submandibular bleed as a baseline, and four hours after gavage *via* terminal cardiac puncture. Plasma was obtained from the supernatant fraction after centrifugation of blood at 2,300 *x g*. Fluorescence of plasma was measured by the EnSight™ Plate Reader (PerkinElmer). All experiments performed were approved by the La Trobe University animal ethics committees (AECs, AEC18024, AEC18036) in accordance with the *Australian code for the care and use of animals for scientific purposes*.

### IEC isolation

Intestinal sections were dissected open, rinsed in DPBS, and incubated in 15 mM EDTA in DPBS for 15 minutes at 37°C with agitation. IECs were dissociated into solution by vortexing. Cells were washed once in chilled DPBS and snap-frozen over dry ice, stored at −80°C until use.

### Western immunoblotting

Cells were lysed on ice for 1 hour in 20 mM Tris pH 7.5, 135 mM NaCl, 1.5 mM MgCl_2_, 1 mM EDTA, 10 % (v/v) glycerol, 1 % (v/v) Triton X-100, supplemented with cOmplete Protease Inhibitor Cocktail (Roche, following the manufacturer’s protocol). Lysates were centrifuged for 5 minutes at 16,000 *x g* and supernatant protein concentrations determined using the Pierce BCA Protein Assay Kit (Thermo Scientific) according to the manufacturer’s protocol. Equalised amounts of protein between 15-40 ng per lane were boiled for 5 minutes at 95°C in reducing buffer (4x stock prepared at 178.3 mM Tris, 350 mM dithiothreitol, 288.4 mM SDS, 670 mM bromophenol blue, 36 % (v/v) glycerol, 10 % (v/v) ϕ3-mercaptoethanol) and electrophoresed on 4-12 % polyacrylamide NuPAGE™, Bis-Tris Mini Protein Gels (Invitrogen) using NuPAGE™ MES SDS Running Buffer (Invitrogen). Proteins were wet transferred in NuPAGE™ Transfer Buffer supplemented with 10 % (v/v) methanol, using the Mini Blot Module (Invitrogen) at 17 V for 1 hour, onto 0.45 μm nitrocellulose membranes (Amersham Protran). Membranes were blocked in 5 % (w/v) skim milk in PBS (2.7 mM KCl, 1.76 mM KH_2_PO_4_, 136.7 mM NaCl, 8.07 mM anhydrous Na_2_HPO_4_) for 1 hour at room temperature (RT) with agitation. Membranes were probed overnight at 4°C with the following antibodies at the following ratios, prepared in 1 % (w/v) skim milk in PBST (PBS + 0.05% (v/v) Tween-20): Beclin1, 1:500 (CST, 3495); ATG7, 1:500 (Sigma-Aldrich, A2856), p62/SQSTM1, 1:500 (CST, 5114); LC3B, 1:500 (Novus Biologicals, NB100-2220); ϕ3-Actin, 1:5000 (Sigma-Aldrich, A2228). Membranes were washed in PBST and probed for 1 hour at RT with the following antibodies, prepared in 1 % (w/v) skim milk in PBST: Donkey anti-Rabbit IgG, 1:10,000 (GE Healthcare, NA943V), Goat anti-Mouse IgG, 1:10,000 (Sigma-Aldrich, A0168). Luminescent signals were visualised using the Western Lightning Plus-ECL (PerkinElmer) kit and ChemiDoc Imaging System (Bio-Rad). Images were processed using Image Lab (Bio-Rad).

### Immunohistochemistry and histology

Formalin-fixed, paraffin embedded intestinal sections of 4 μm thickness were dewaxed with xylene and rehydrated into RO water *via* decreasing ethanol grades. For immunohistochemistry, antigen retrieval was performed by boiling slides in citrate buffer (10 mM sodium citrate, 0.05 % (v/v) Tween-20) for 20 minutes. Slides were quenched for 20 minutes in 3 % (w/w) hydrogen peroxidase and rinsed in TBST (0.5 M Tris, 9 % (w/v) NaCl, 0.5 % (v/v) Tween-20, pH 7.6). Sections were incubated overnight in a humidified chamber with primary antibodies diluted in SignalStain® Antibody Diluent (Cell Signalling Technology (CST), 8112) at the following ratios: Cleaved caspase-3, 1:200 (CST, 9661); Ki-67, 1:150 (Invitrogen, MA5-14520); Lysozyme, 1:300 (Thermo Scientific, RB-372-A1); Chromogranin A, 1:500 (Abcam, ab85554); DCAMKL1, 1:400 (Abcam, ab31704). Slides were washed in TBST and antigens detected using the EnVision+ System (DAKO, K4003) following the manufacturer’s protocol. Slides were counterstained with Mayer’s haematoxylin, dehydrated in increasing ethanol grades, cleared in xylene and mounted with DPX (Sigma-Aldrich). For TUNEL staining, the TUNEL Assay Kit (Abcam, ab206386) was used according to the manufacturer’s protocol. For IAP staining, slides were dewaxed and rehydrated into PBST (see Western immunoblotting), and the Vector Red Alkaline Phosphatase Substrate Kit (Vector Laboratories SK-5100) was used according to the manufacturer’s protocol and included counterstaining with Mayer’s haematoxylin. For PAS-AB staining, dewaxed and rehydrated slides were incubated for 15 minutes in 1 % (v/v) Alcian blue in 3 % (v/v) aqueous acetic acid (Amber Scientific), 5 minutes in 1% (v/v) aqueous period acid (Amber Scientific), and 10 minutes in Schiff’s reagent (Amber Scientific, with RO water rinses in between steps. Slides were also counterstained with Mayer’s haematoxylin.

### Transmission electron microscopy

Duodenal tissue was fixed in Karnovsky’s fixative (2.5 % (v/v) glutaraldehyde, 2 % (v/v) paraformaldehyde, 0.1 M sodium cacodylate) for 4 hours at 4°C. Tissue was post-fixed in 1 % (w/v) osmium tetroxide, 1.5 % (w/v) potassium ferrocyanide and 0.1 M sodium cacodylate, for 3 hours at RT. Tissues were rinsed in Milli-Q water and dehydrated in increasing grades of ethanol. Samples were embedded into Spurrs resin by two immersions in 100 % acetone for 20 minutes each, then two hour immersions in mixtures of Spurrs to acetone at 1:3, 1:1 and 3:1 ratios. Overnight infiltration in 100 % Spurrs was followed by 2 hours in 100 % Spurrs under vacuum conditions. Blocks were polymerised overnight at 70°C. 70 nm ultrathin sections were contrasted for 10 minutes in 10 % (w/v) uranyl acetate in methanol under dark conditions, rinsed sequentially with 50 % (v/v) methanol, 25 % (v/v) methanol then Milli-Q water, followed by incubation in Reynold’s lead citrate solution (0.11 M lead nitrate, 0.19 M sodium citrate, pH 12, prepared in CO_2_-free Milli-Q water) for 10 minutes in carbon dioxide depleted conditions. Sections were mounted onto a copper grid, carbon-coated, and imaged with the Joel JEM-2100 transmission electron microscope at 80kV equipped with Gatan digital camera.

### Organoid culture and cell death assays

Organoids were established by culturing crypt-enriched fractions from the duodenum of untreated mice. To isolate crypt-enriched fractions, duodenal tracts were opened and rinsed in DPBS. Villi were removed by gentle scraping. Tracts were cut into 0.5 cm sections, rinsed vigorously with DPBS, and incubated in 2 mM EDTA in DPBS for 30 minutes at 4°C with gentle agitation. Sections were washed once in cold DPBS and crypts obtained by dissociation from the stromal tissue into DPBS by vigorous pipetting. Crypts were resuspended in Advanced DMEM media (Advanced DMEM (Gibco), 1 % (w/v) bovine serum albumin, 20 mM L-Glutamine, 100 mM HEPES, 1000 U/mL Penicilln-Streptomycin) and passed through a 70 μm strainer. Remaining single cells were excluded by repeat low speed column centrifugation (76 *x g*, 2 minutes) and discarding of cloudy supernatant. Crypt-enriched pellets were then resuspended in Cultrex Reduced Growth Factor Basement Membrane Extract, Type 2, Path Clear (R&D Systems) at 50-300 crypts per 50 μL dome in pre-warmed 24 well plates. Domes were polymerised by incubation for 15 minutes at 37°C and Advanced DMEM media supplemented with 1x N-2 (Gibco), 1x B-27 (Gibco), 0.05 ng/μL human EGF (Peprotech, AF-100-15), 0.1 ng/μL murine noggin (Peprotech, 250-38), 0.5 ng/μL murine R-Spondin-1 (Peprotech, 315-32) was added. Organoids were maintained at 37°C, 5 % CO_2_, with media being replaced every 2-3 days. Passaging was performed every 7-10 days by mechanically dissociating organoids and re-seeding into fresh Cultrex at a 1:3 to 1:6 split. To induce deletion, organoids were seeded into media containing 200 nM 4-HT (Sigma-Aldrich, H7904) for three days, and then maintained as per normal. For apoptosis assays, organoids were dissociated by incubation for 15 minutes at 37°C in TrypLE Express (Gibco) plus pipetting. Cells were stained in the dark for 15 minutes with either FITC Annexin V (BD Pharmingen, 556419) or APC Annexin V (BD Pharmingen, 550475) diluted at 1:20, and Propidium iodide solution (Sigma-Aldrich, P4864) diluted at 1:200 in 1x Annexin V Binding Buffer (BD Pharmingen, 556454). Stained cells were diluted 4x in 1x Annexin V Binding Buffer and analysed by flow cytometry performed using the BD FACSCanto II (Becton Dickinnson and Company, BD Biosciences, San Jose, CA, USA). Data was analysed using FlowJo^TM^ Software, Version 10.5.3 (Becton Dickinnson and Company, BD Biosciences, San Jose, CA, USA).

### Intestinal organoid whole-mount immunofluorescence

Whole-mount intestinal organoid staining protocol was adapted from Dekkers *et al* ^41^. Briefly, organoids were removed from matrix by incubation with ice-cold Gentle Cell Dissociation Reagent (STEMCELL Technologies, 100-0485) with gentle rocking at 4°C for 60 minutes. Organoids were then fixed in 4 % (w/v) paraformaldehyde solution (ProSciTech, Catalog no. C004) and blocked with Organoid Wash Buffer (dPBS (Gibco^TM^, 14190144) + 0.1% (w/v) Triton X-100 (Merck, X100) + 0.2 % (w/v) Bovine Serum Albumin (Merck, A3059)), followed by incubation with rat anti-E-cadherin primary monoclonal antibody ECCD-2 (Invitrogen, 13-1900) diluted at 1:500 overnight at 4°C with gentle rocking. Following primary antibody incubation, organoids were washed extensively and incubated overnight with a combination of goat anti-Rat IgG (H+L) cross-adsorbed secondary antibody, Alexa Fluor^TM^ 568 (Invitrogen, A-11077) at 1:500 dilution and nuclear stain DAPI (1 μg/ml, Merck, D9542) at 4°C with gentle rocking. Organoids were then subjected to another extensive washing step prior to sample clearing and mounting steps. In lieu of the fructose-glycerol solution used for organoid clearing and mounting as outlined in Dekkers *et al* ^41^, RapiClear® 1.49 solution (SunJin Lab, RC149001) was used instead in accordance with the manufacturer’s protocol. Images were then acquired using Zeiss LSM 980 with Airyscan 2 confocal microscope. Airyscan 2 Multiplex SR-4Y imaging mode was utilised and parameters such as laser power and gain were kept consistent amongst samples. Images were taken using the 40x (water) objective with 1.7x Zoom and imaged once to avoid excessive photobleaching. Images were then analysed using Zen Blue (Zeiss) and ImageJ software (Fiji ^42^). 3D rendering video was created using “3D Project” function in ImageJ.

### Intraepithelial lymphocytes isolation for flow cytometry analysis

Intraepithelial lymphocytes were isolated from the small intestines and colons of mice as previously described. Briefly, excess fat was removed from the colon and small intestines. Peyer’s patches were also removed from the small intestines. The tissues were kept moist by frequent saturation with ice-cold wash buffer (dPBS + 2 % (w/v) fetal calf serum). Intestines were cut longitudinally through the lumen, and fecal matter washed away by lightly swirling in a petri dish containing ice-cold wash buffer. Intestines were then cut into 0.5 cm fragments and placed in dissociation solution (wash buffer + 5 mM EDTA) and incubated for 30 minutes at 37°C with gentle shaking. The cell suspension was then vortexed and filtered using a 70 μm cell strainer, transferred to a new 50 mL tube and centrifuged at 1700 rpm for 7 minutes. The cell pellets were then washed twice with wash buffer. Cells were then resuspended in discontinuous 40 % / 80 % Percoll® (GE Healthcare, 17089101) gradient and centrifuged at 900 x *g* for 20 minutes at room temperature. The interface was collected and washed twice with wash buffer and the final pellet containing intraepithelial lymphocytes stained.

For flow cytometry analysis, isolated IELs from above were incubated with Live/Dead marker (1:500, BD Biosciences, 566332), anti-CD45.2 (1:200, BD Biosciences, 741957), CD3ε (1:200, BD Biosciences, 564378), CD4 (1:1000, BD Biosciences, 741461), CD8α (1:1000, BD Biosciences, 562283), CD8β (1:1000, BD Biosciences, 741127), TCRβ (1:300, ThermoFisher Scientific, 47-5961-82), TCRγδ (1:300, BD Biosciences, 563994), NK1.1 (1:1000, BD Biosciences, 566502), FOXP3 (1:300, Thermofisher Scientific, 12-5773-82), and CD19 (1:300, BD Biosciences, 563333) for total IEL analysis. For intracellular staining, cells were fixed and permeabilised using the Foxp3/Transcription factor staining buffer kit (Invitrogen, 00-5523-00) according to the manufacturer’s instructions, followed by incubation with intracellular markers. Finally, cells were resuspended in FACS wash buffer containing CountBright Absolute Counting Beads (Molecular Probes, C36950) and stained cell acquisition was performed on the BD FACSymphony^TM^ A3 Cell Analyzer (Becton Dickinnson and Company, BD Biosciences, San Jose, CA, USA) and data analysed using FlowJo^TM^ Software, Version 10.5.3 (Becton Dickinnson and Company, BD Biosciences, San Jose, CA, USA).

## Acknowledgements

We thank Prof. Joan Heath and Dr. Karen Doggett for providing the UBC-Cre-ERT2 mice (Walter and Eliza Hall Institute, Melbourne, VIC, Australia). We acknowledge scholarship support for S.T. (La Trobe University Research Training Program Scholarship), L.J.J. (La Trobe University Australian Postgraduate Award) and J.J. (La Trobe Graduate Research Scholarship and Full Fee Research Scholarship). We are grateful to the Australian Research Council for grant support (E.F.L., W.D.F., J.M.M., DP190102612), and to the National Health and Medical Research Council (J.M.M., 1046092; K.D., A.S.Y., 2010704, 1136592), the Victorian Cancer Agency (E.F.L., MCRF19045) for fellowship support.

## Author contributions

S.T., J.J. designed, performed, and analyzed experiments and wrote the paper; T.J.H., M.E., J.R. S.L.E., C.M.R. R.N., I.Y.L., L.J.J., S.G., M.Y., C.I., D.B. designed, performed experiments, analyzed and interpreted data; M.B., J.C.L., P.D.C., K.D., P.A.G., B.T.K., L.A.M., A.S.Y., J.M.M. analyzed and interpreted data; W.D.F., E.F.L. designed the project, analyzed, interpreted data, and wrote the paper. All authors commented on the manuscript.

## Competing interests

All authors declare no competing interests.

## Additional information

Correspondence and requests for materials should be addressed to W. Douglas Fairlie or Erinna F. Lee.

---

**Supplementary Fig. 1 PCR-based genotyping following tamoxifen-induced deletion of *Becn1* and *Atg7* in adult *Becn1^fl/fl^*;*Vil1-CreERT2^Cre/+^*- and *Atg7^fl/fl^*;*Vil1-CreERT2^Cre/+^* - derived intestinal epithelial cells.** Stom: stomach. Duo: duodenum. Jej: jejunum. Ile: ileum. Col: colon. KO: knock-out. WT: wild-type

**Supplementary Fig. 2 Intestinal epithelium-specific loss of ATG7 in adult mice leads to slightly longer small intestinal lengths and decreased body weight gain which develop over one month. a** PCR-based genotyping and **b** Western blotting demonstrated successful deletion of ATG7 in intestinal epithelial cells. Representative images of **c** abdominal necropsy and **d** intestinal tracts demonstrating slightly increased small intestinal, but not colon, lengths. However, there was no evidence of severe loss of intestinal homeostasis seen in the absence of BECLIN1 over a significantly shorter time frame. **e** The absence of ATG7 over this extended period of time leads to a decrease in body weight gain. All graphs show the mean ± S.E.M. Significance was determined by Wilcoxon’s t-test. Stom: stomach. Duo: duodenum. Jej: jejunum. Ile: ileum. Col: colon. KO: knock-out. WT: wild-type. *: non-specific band.

**Supplementary Fig. 3 Immunophenotyping of the intraepithelial lymphocytes in the small intestines.** Graphs show the frequency of each indicated immune cell subtype within total live cells. Data represent mean ± S.E.M and significance determined by ordinary one-way ANOVA. Data is representative of two independent experiments.

**Supplementary Fig. 4 Tamoxifen-induced deletion of *Becn1* and *Atg7* in *Becn1^fl/fl^*;*Vil1-CreERT2^Cre/+^*- and *Atg7^fl/fl^*;*Vil1-CreERT2^Cre/+^* -derived intestinal organoids as detected by a PCR genotyping and b Western blotting.** *: non-specific band.

## Notes

### Competing Interest Statement

The authors have declared no competing interest.

