## Supplemental Material for "BECLIN1 is essential for intestinal homeostasis"

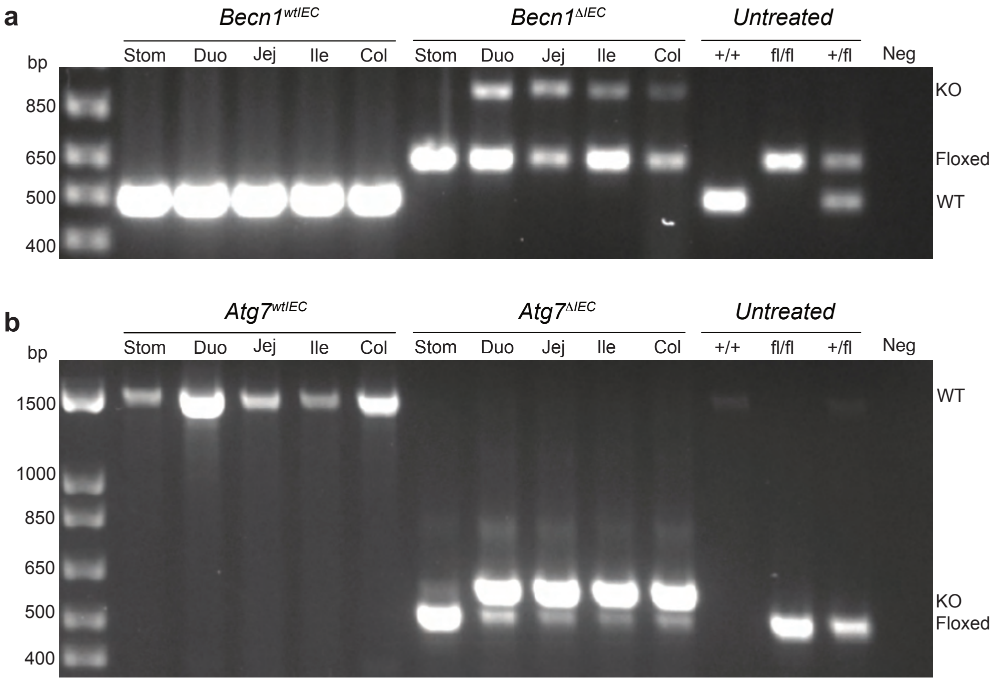


**Supplementary Fig. 1 PCR-based genotyping following tamoxifen-induced deletion of *Becn1* and *Atg7* in adult *Becn1*$^{fl/fl}$;*Vil1-CreERT2*$^{Cre/+}$- and *Atg7*$^{fl/fl}$;*Vil1-CreERT2*$^{Cre/+}$ -derived intestinal epithelial cells.** Stom: stomach. Duo: duodenum. Jej: jejunum. Ile: ileum. Col: colon. KO: knock-out. WT: wild-type.

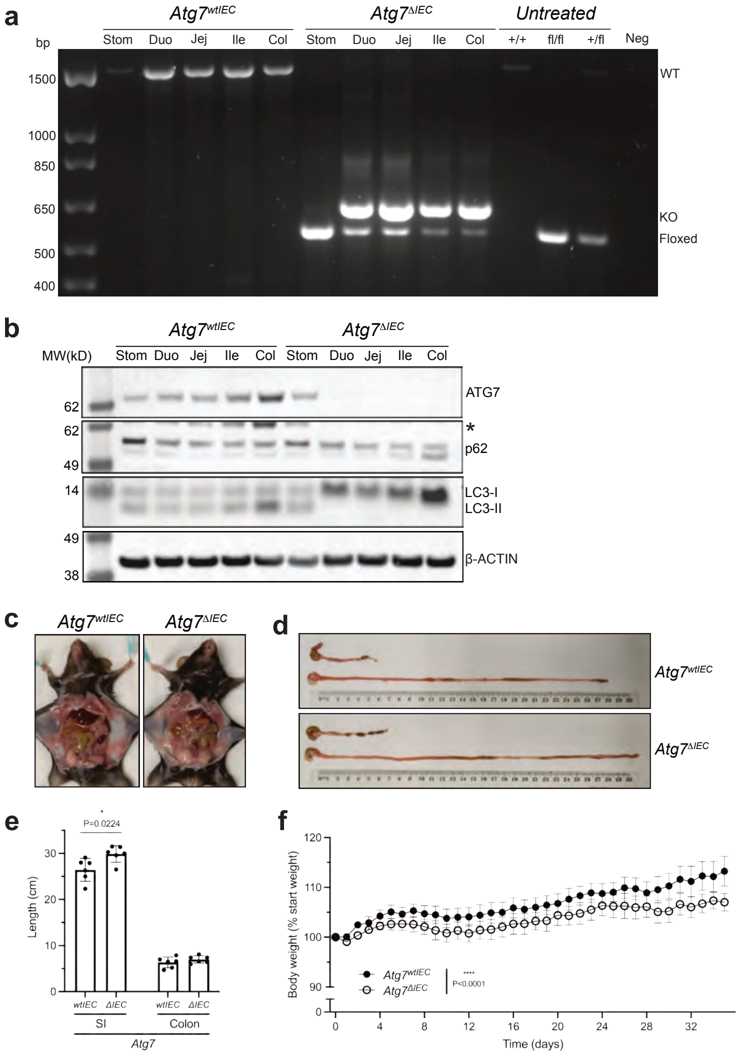


**Supplementary Fig. 2 Intestinal epithelium-specific loss of ATG7 in adult mice leads to slightly longer small intestinal lengths and decreased body weight gain which develop over one month. a** PCR-based genotyping and **b** Western blotting demonstrated successful deletion of ATG7 in intestinal epithelial cells. Representative images of **c** abdominal necropsy and **d** intestinal tracts demonstrating slightly increased small intestinal, but not colon, lengths. However, there was no evidence of severe loss of intestinal homeostasis seen in the absence of BECLIN1 over a significantly shorter time frame. **e** The absence of ATG7 over this extended period of time leads to a decrease in body weight gain. All graphs show the mean ± S.E.M. Significance was determined by Wilcoxon's t-test. Stom: stomach. Duo: duodenum. Jej: jejunum. Ile: ileum. Col: colon. KO: knock-out. WT: wild-type. *: non-specific band.

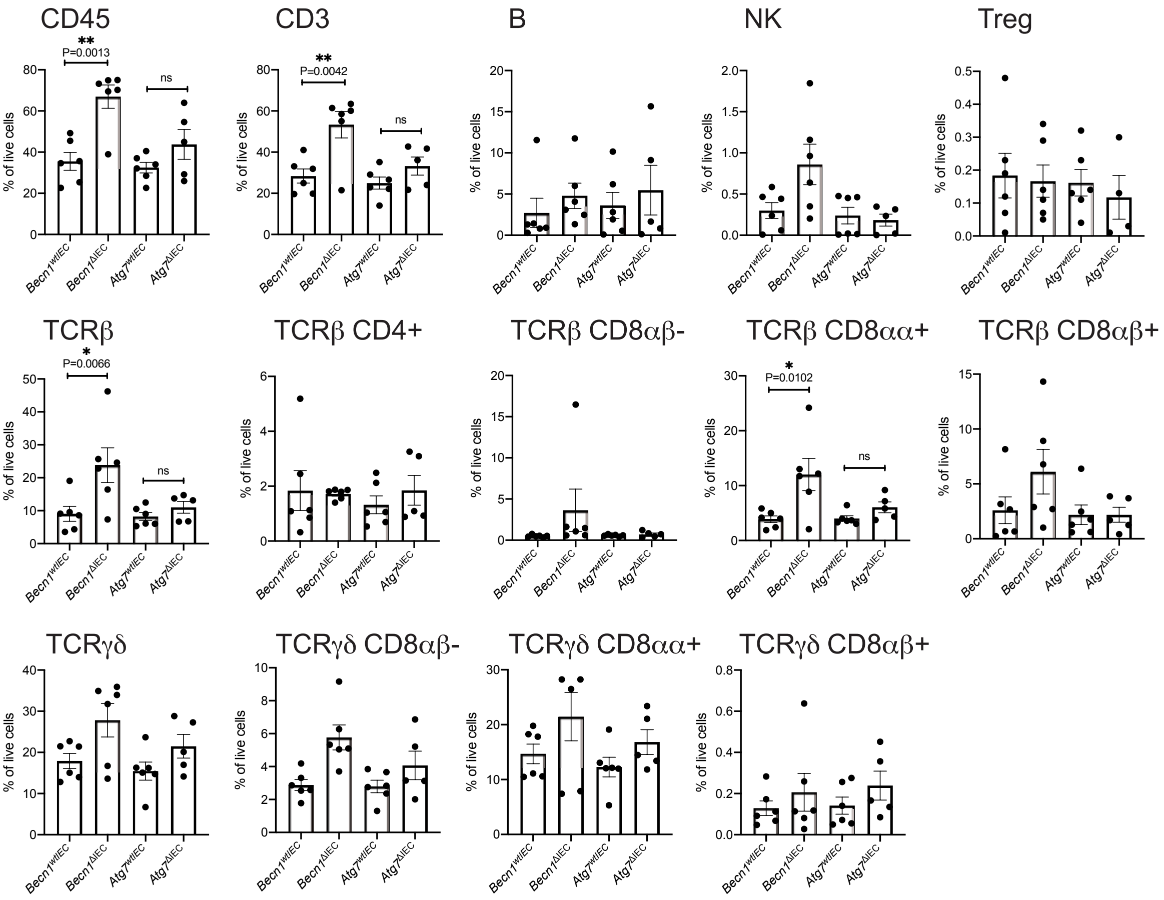


**Supplementary Fig. 3 Immunophenotyping of the intraepithelial lymphocytes in the small intestines.** Graphs show the frequency of each indicated immune cell subtype within total live cells. Data represent mean ± S.E.M and significance determined by ordinary one-way ANOVA. Data is representative of two independent experiments.

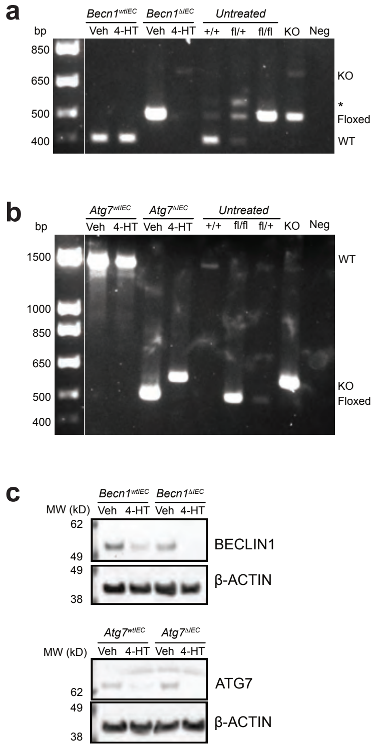


**Supplementary Fig. 4 Tamoxifen-induced deletion of *Becn1* and *Atg7* in *Becn1*$^{fl/fl}$;*Vil1-CreERT2*$^{Cre/+}$- and *Atg7*$^{fl/fl}$;*Vil1-CreERT2*$^{Cre/+}$ -derived intestinal organoids as detected by a PCR genotyping and b Western blotting.** *: non-specific band.
